# Comparative genomics of *Mycobacterium africanum* Lineage 5 and Lineage 6 from Ghana suggests different ecological niches

**DOI:** 10.1101/202234

**Authors:** Isaac Darko Otchere, Mireia Coscollá, Leonor Sánchez-Busó, Adwoa Asante-Poku, Daniela Brites, Chloe Loiseau, Conor Meehan, Stephen Osei-Wusu, Audrey Forson, Clement Laryea, Abdallah Iddrisu Yahayah, Akosua Baddoo, Gloria Akosua Ansa, Samuel Yaw Aboagye, Prince Asare, Sonia Borrell, Florian Gehre, Patrick Beckert, Thomas A Kohl, Sanoussi N'dira, Christian Beisel, Martin Antonio, Stefan Niemann, Bouke C de Jong, Julian Parkhill, Simon R Harris, Sebastien Gagneux, Dorothy Yeboah-Manu

## Abstract

*Mycobacterium africanum* (*Maf*) causes up to half of human tuberculosis in West Africa, but little is known on this pathogen. We compared the genomes of 253 *Maf* clinical isolates from Ghana, including both L5 and L6. We found that the genomic diversity of L6 was higher than in L5, and the selection pressures differed between both groups. Regulatory proteins appeared to evolve neutrally in L5 but under purifying selection in L6. Conversely, human T cell epitopes were under purifying selection in L5, but under positive selection in L6. Although only 10% of the T cell epitopes were variable, mutations were mostly lineage-specific. Our findings indicate that *Maf* L5 and L6 are genomically distinct, possibly reflecting different ecological niches.

## Introduction

The global phylogeography of the human-adapted *Mycobacterium tuberculosis* complex (MTBC) demonstrates highest diversity in West Africa, with six out of the seven known lineages represented^1,2^. Two of these lineages, Lineage 5 (L5) and Lineage 6 (L6), together originally known as *Mycobacterium africanum* (*Maf*), are restricted to West Africa for unknown reasons. By contrast, MTBC lineages belonging to *Mycobacterium tuberculosis* sensu stricto (*Mtbss*), in particular Lineage 4 (L4), are more geographically widespread^1^. *M. africanum* has remained an important pathogen in West Africa since its first description in 1968^3^, and is responsible for up to half of human tuberculosis (TB) in some regions^4^.

The MTBC is thought to have originally emerged in Africa and subsequently spread to other parts of the world following waves of human migrations, trade and conquests^5–8^. Yet the reason(s) why *Maf* is limited to West Africa despite, for example, centuries of the trans-Atlantic slave trade remains unknown. Some comparative studies have identified phenotypic differences between the two *Maf* lineage^9,10^, suggesting they might be fundamentally distinct and occupy different ecological niches.

Three hypotheses have been put forward to explain the restriction of *Maf* to West Africa. The first hypothesis proposes that *Maf* might have emigrated outside of Africa but was later outcompeted by *Mtbss*, which has been shown to be more virulent than *Maf* in animal models^11^. The second hypothesis states that the restriction of *Maf* to West Africa is due to its adaptation to West African human populations^9,12^. Finally, according to the third hypothesis, *Maf* might be zoonotic with an animal reservoir restricted to West Africa.

Some evidence in support of the first hypothesis is the reported association of *Maf* (L6) with HIV co-infection, attenuated ESAT-6 responses and delayed progression to active disease relative to *Mtbss*^9,13–16^. In addition, both *Maf* lineages as well as *Mtbss* L1, together described as “ancestral” MTBC lineages, have been shown to elicit a stronger early production of pro-inflammatory cytokines compared to the “modern” MTBBC L2, L3 and L4^17^. The delayed pro-inflammatory immune response in the “modern” MTBC lineages might allow for more rapid disease progression and transmission^17^. The second hypothesis is supported by the statistical association of L5 with the native West African ethnic group known as “Ewe” reported by two independent studies in Ghana^9,12^. The third hypothesis is mainly supported by the phylogenetic placement of *Maf* (L6) amidst the cluster of the animal-adapted members of the MTBC in the various phylogenies of the MTBC^5,7,18^.

If the first hypothesis is true, the proportion of *Maf* associated TB in West Africa is expected to decline over time. However, there are conflicting reports of the proportion of *Maf* associated TB in West Africa. Even though the report of a steady decline of *Maf* associated TB in some settings seems to support the first hypothesis^19–21^, other studies indicate that *Maf* remains an important cause of TB in West-Africa^22–24^. In Ghana for instance, a recent study showed that the proportion of TB due to *Maf* remained constant over the 8 year study period^25^. Even though, the reported statistical association of L5 with ethnicity in Ghana suggests a possible co-evolutionary scenario in favour of the second hypothesis, genetic evidence of co-evolution/co-adaptation remains to be demonstrated. In the case of the third hypothesis, the environmental or zoonotic reservoir(s) need to be identified.

In this study, we used whole genome sequencing of Ghanaian *Maf* clinical strains to explore genomic differences between the two *Maf* lineages that might support one or more of these hypotheses.

## Results

### Whole genome SNP distance, average nucleotide diversity and phylogeny of *Maf* in Ghana

Our data set comprised *Maf* isolates obtained from TB patients reporting to various hospitals in Ghana. After excluding genomes that did not meet the criteria for mapping (Supplementary figure S1), 253 *Maf* genomes (175 L5 and 78 L6) were used for the analysis. Patients’ residential regions are provided (Supplementary figure S2). The upper right pie chart indicates those 97 patients (55 infected with L5 and 42 with L6) with no information on region of residence. We found the number of fixed SNPs (SNPs found in more than 95% of genomes) in a genome compared to the MTBC ancestor^26^ to be significantly higher in L6 (1,037) compared to L5 (928) (Wilcoxon rank-sum test, p < 0.0001) (Fig 1A). Moreover, despite the larger number of L5 genomes (more than twice the number of L6 genomes) analyzed, the mean pairwise SNP distance between any two strains was significantly higher in L6 (360) compared to L5 (223) (Wilcoxon rank-sum test, p < 0.0001; Fig 1B). Finally, the whole genome average nucleotide diversity (*π*) for L6 (0.000110) was significantly higher compared to L5 (0.00007) (Fig 1C, non-overlapping 95% confidence interval (CI)). Taken together, these findings show that L6 in Ghana is significantly more genetically diverse than L5 irrespective of sample size. The whole genome-based phylogenetic tree of the Ghanaian *Maf* strains generated from 11,027 total polymorphic positions between the *Maf* strains and the MTBC ancestor reference excluding repetitive and mobile genetic element rooted on *M. canettii* is shown in Figure 2. The *Maf* lineages were resolved as two distinct branches of the genome-based tree with possible sub-groups (Fig 2).

**Figure 1:**
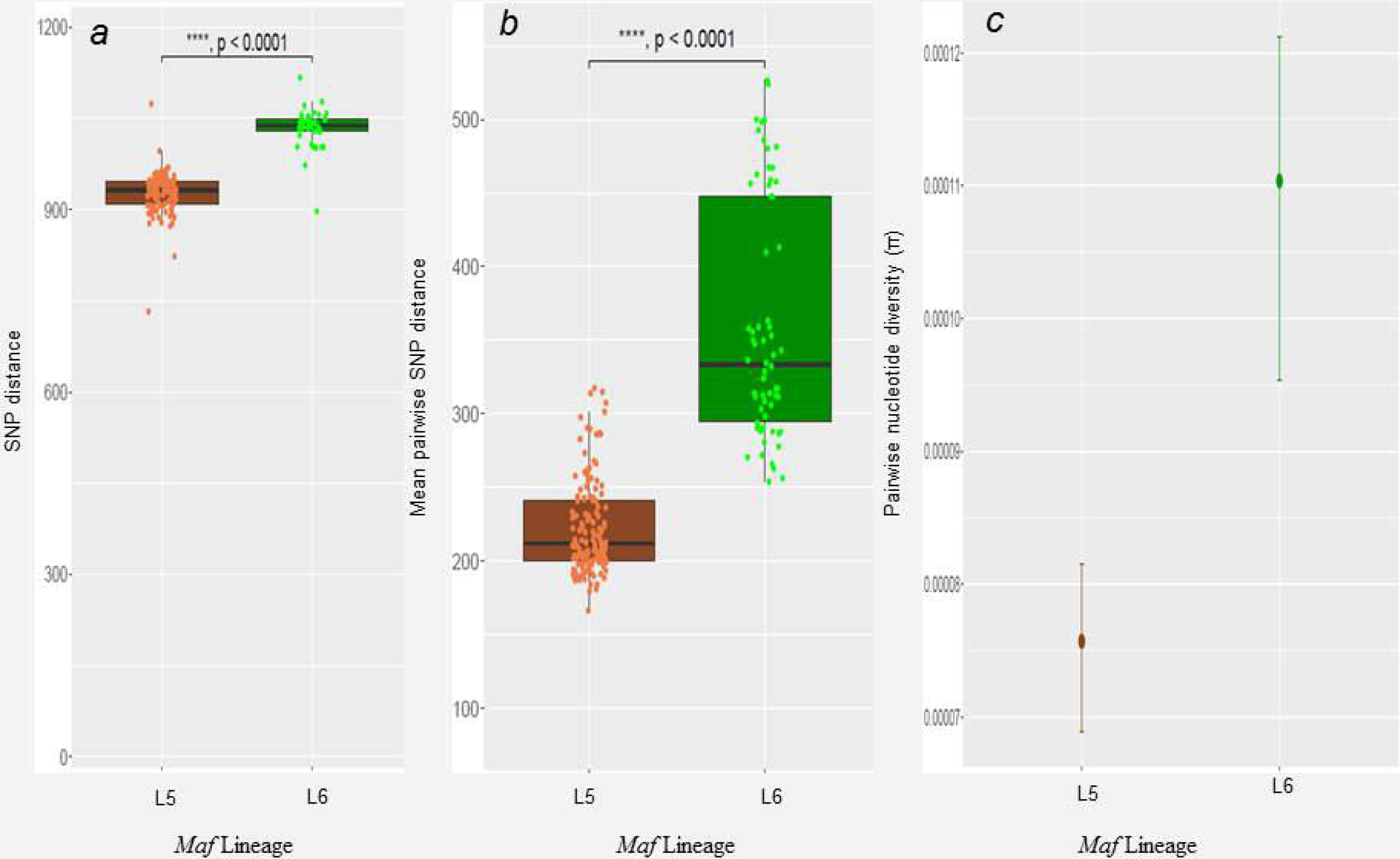
Number of SNPs per *Maf* Lineage (175 L5 and 78 L6 genomes). *a*: Number of SNPs between *Maf* genomes and the hypothetical MTBC ancestor (the median fixed SNPs of L5 (934) is lower (W = 417, p-value < 2.2e-16) compared to L6 (1,039). *b*: Pairwise SNPs between genomes within each lineage (the median of the pairwise SNPs is lower (W = 234, p-value < 2.2e-16) in L5 (212) compared to L6 (334). *c*: Whole genome average nucleotide diversity (n) between L5 and L6 (the mean diversity of L5 (0.000076) is significantly (non-overlapping 95% confidence intervals) lower than L6 (0.000110). Error bars indicate 95% confidence intervals.

**Figure 2:**
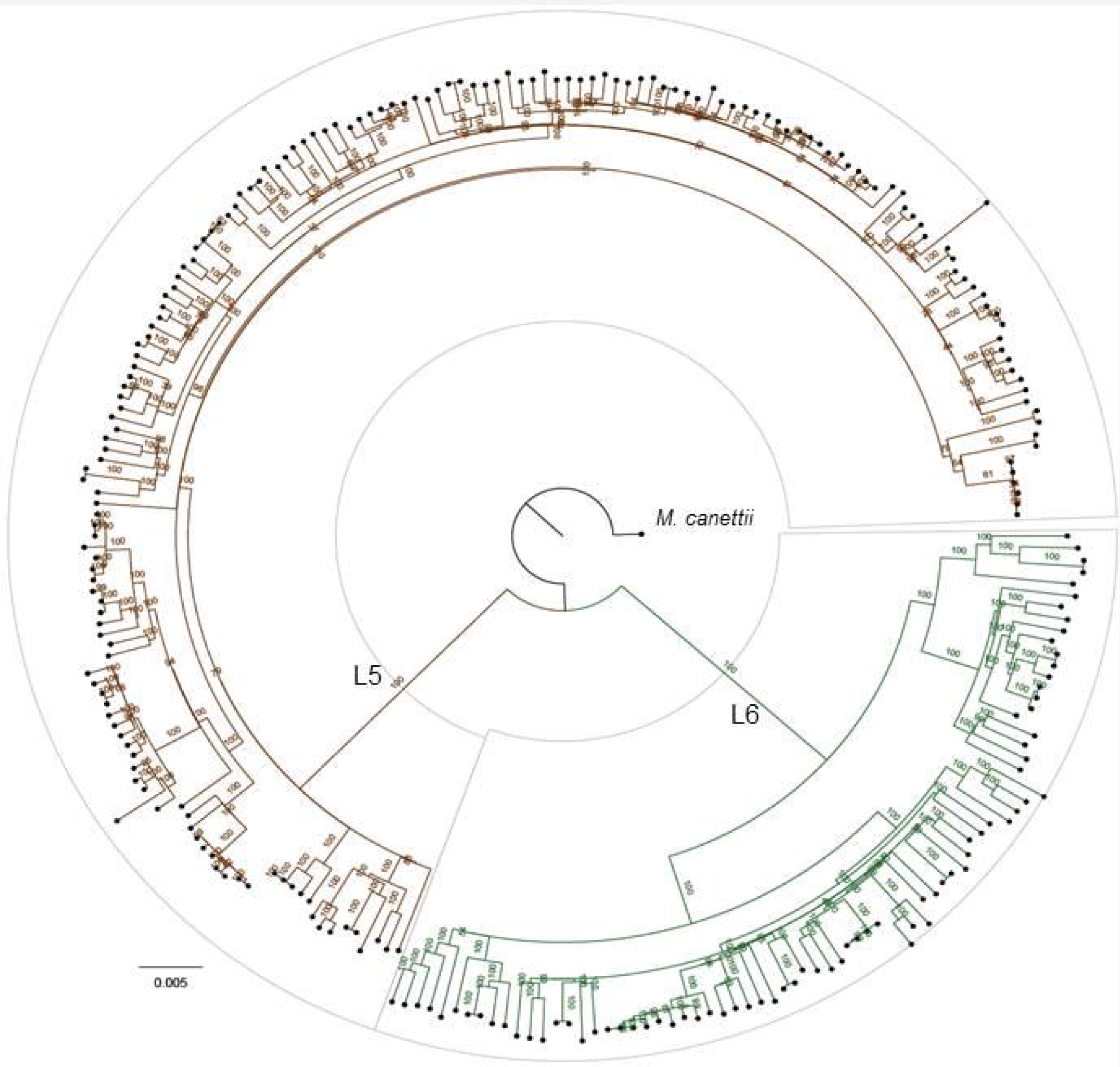
Phylogeny of Ghanaian *Maf* strains. (The maximum likelihood phylogenetic tree of 253 Ghanaian *Maf* isolates is based on 11,027 variable positions. The tree was rooted on *M. canettii* and the confidence of nodes was assessed by bootstrapping 1000 pseudo replicates. Each lineage clade is colored according to the conventional MTBC lineage color codes^1^.

### Genetic diversity of L6 is significantly higher than L5 among T cell epitopes and genes of other functional categories

We found that the higher diversity of L6 compared to L5 was reflected across all the 8 functional categories of genes analyzed (Fig. 3). Whereas pairwise nucleotide diversity (*π*) for L5 was below 0.0001 across all functional categories, the estimates for L6 were all above 0.0001. The most prominent difference between L6 and L5 was within 1,226 experimentally confirmed human T cell epitopes of MTBC which we downloaded from the Immune Epitope Database (IEDB)^27^, for which the mean *π* for L5 was 0.000063 compared to the 0.000149 estimated for L6, reflecting more than a two-fold difference in diversity (non-overlapping 95% CI).

**Figure 3.**
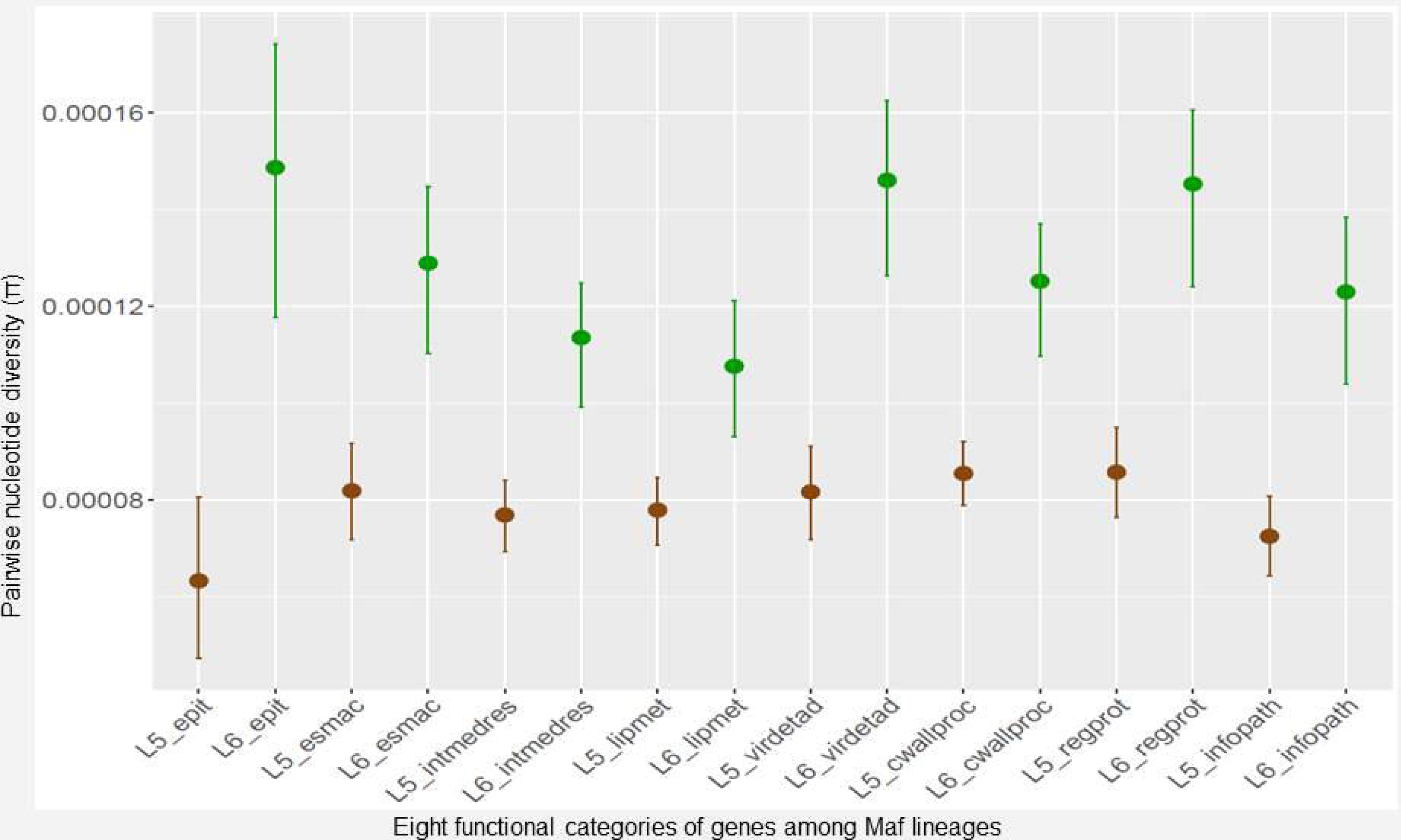
Averaged nucleotide diversity (*π*) of *Maf* within genes of eight functional categories. *epit* - genes encoding human T cell epitopes, *esmac* - genes essential for growth in macrophages, *intmedres* - genes involved with intermediate metabolism and respiration, *lipmet* - genes involved with lipid metabolism, *virdetad* - genes involved with virulence, detoxification and adaptation, *cwallproc* - genes involed with cell wall and cell processes, *regprot* - genes encoding regulatory proteins and *infopath* - genes involved with information pathways. Error bars are indications of 95% confidence intervals.

Within L5, there was no difference between the estimated *π* for the T cell epitopes and any of the other functionally categorized genes. However, within L6, genes encoding regulatory proteins and those involved with virulence, detoxification and adaptation were more diverse compared to those for lipid metabolism as well as intermediate metabolism and respiration (non-overlapping 95% CI). In addition, genes encoding regulatory proteins were more diverse compared to those involved with lipid metabolism (non-overlapping 95% CI).

### Different selection pressures within L5 and L6 in human T cell epitopes and regulatory proteins

The average pairwise dN/dS of the concatenates of T cell epitopes as well as genes of the seven other functional categories were calculated for all genomes and compared between L5 and L6. Apart from sequences encoding human T cell epitopes and regulatory proteins that had median average pairwise dN/dS ratios greater than 1.0 in L5 and L6, respectively (Fig 4, panel a and b), all the remaining functional categories showed a dN/dS ratio of less than 1.0 in both lineages (Supplementary figure S3). Human T cell epitopes of *Maf* L5 (median pairwise dN/dS = 0.64) were significantly more conserved compared to L6, which exhibited higher diversity (median pairwise dN/dS = 1.53) (Wilcoxon rank-sum test, W = 2265, *p* < 0.0001). Conversely, genes encoding regulatory proteins were more diverse among L5 genomes (with median pairwise dN/dS = 1.03) (Wilcoxon rank-sum test, W = 6303, *p* = 0.0010) compared to L6 (with median pairwise dN/dS = 0. 85). To account for the different sample sizes; 147 L5 compared to 67 L6 genomes after excluding 43 genomes differing from others with less than 10 SNPs difference (see Methods and supplementary figure S1), we repeated the analysis using mean values of 10 randomly sampled sets of L5 genomes with sample size 67 among human T cell epitopes (Fig 4, panel c) and regulatory proteins (Fig 4 panel d) and got similar results (Wilcoxon rank-sum test, W = 1300, *p* < 0.0001, W = 3466, *p* < 0.0001 for T cell epitopes and regulatory proteins, respectively).

**Figure 4:**
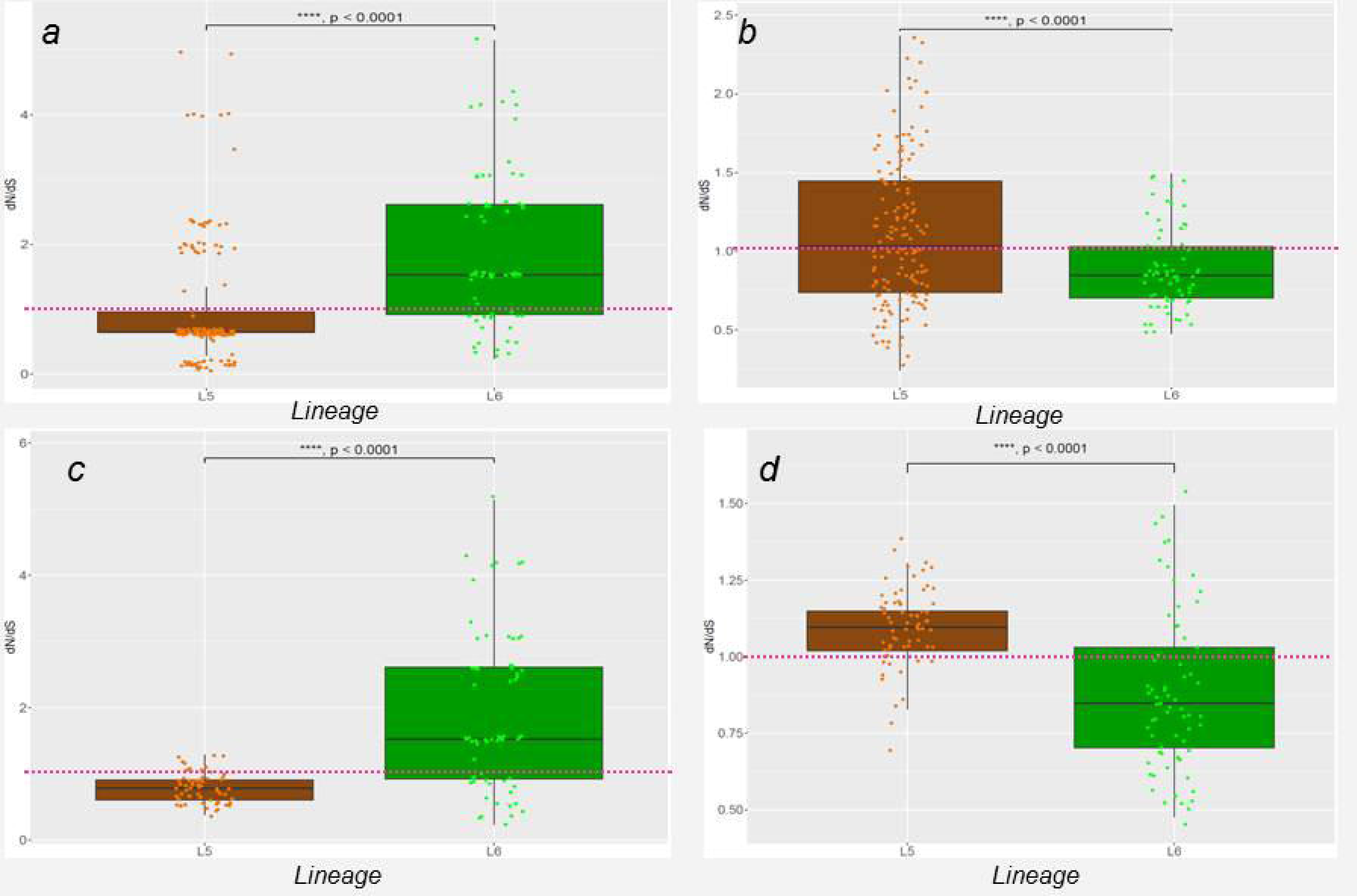
Pairwise dN/dS of genes encoding human T cell epitopes and regulatory proteins in L5 and L6. Estimation of pairwise dN/dS of epitopes (*a*) and regulatory proteins (*b*) using the entire 147 L5 against the 67 L6 genomes. Estimation of pairwise dN/dS of epitopes (*c*) and regulatory proteins (*d*) using the mean dN/dS values of 10 random samples (size =67, with replacement) of L5 against the 67 L6 genomes.

### Lineage-specific accumulation of mutations within human T cell epitopes

When we compared the number of epitopes with amino acid mutations between lineages, we found more epitopes mutated in L6 (57) compared to L5 (45), but no statistically significant difference (Fig 5). In addition, we compared the number of nonsynonymous polymorphic sites between the two Maf lineages within the human T cell epitopes (Supplementary Fig S4), and found that these were more frequent in L6 (38) compared to L5 (28) but with no statistically significant difference between L5 and L6. We compared the identity of the mutant human T cell epitopes between the two *Maf* lineages (Fig 6A) and found 72 epitopes that were uniquely mutated in L5 (among 174 genomes) compared to 54 epitopes in L6 (among 67 genomes). Only two epitopes (IEDB IDs 178644 and 178609) were mutated in both lineages. However, the mutations were at different loci with different amino acid substitutions (A183G and G278D in L5 compared to A177V and D277N in L6). In terms of T cell antigens, there were 28 uniquely mutated in L5 compared to 19 in L6 and 12 mutated in both lineages involving different epitopes within the respective antigens (Fig 6B). The 12 T cell antigens mutated in both lineages are summarized in Table 1. All T cell epitopes and antigens with mutations among the two *Maf* lineages are listed in Supplementary table S5.

**Figure 5:**
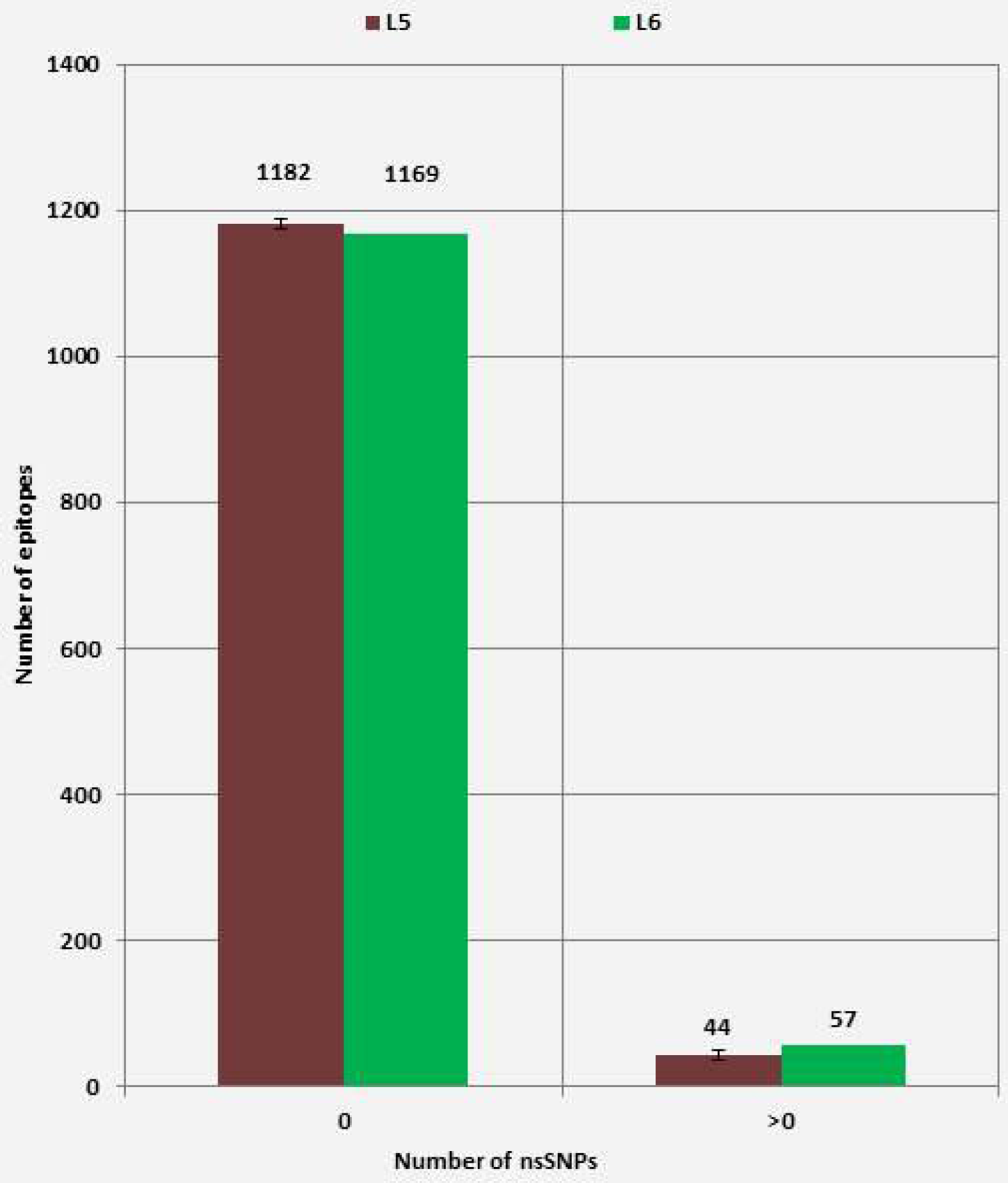
Number of human T cell epitopes with nonsynonymous SNPs (nsSNPs) stratified by *Maf* lineage. No significant difference (X-squared = 1.487, df = 1, p-value = 0.22) between the number of epitopes with nsSNPs among the 67 L6 genomes and L5 (mean values of 10 random samples of size=67 with replacement).

**Figure 6:**
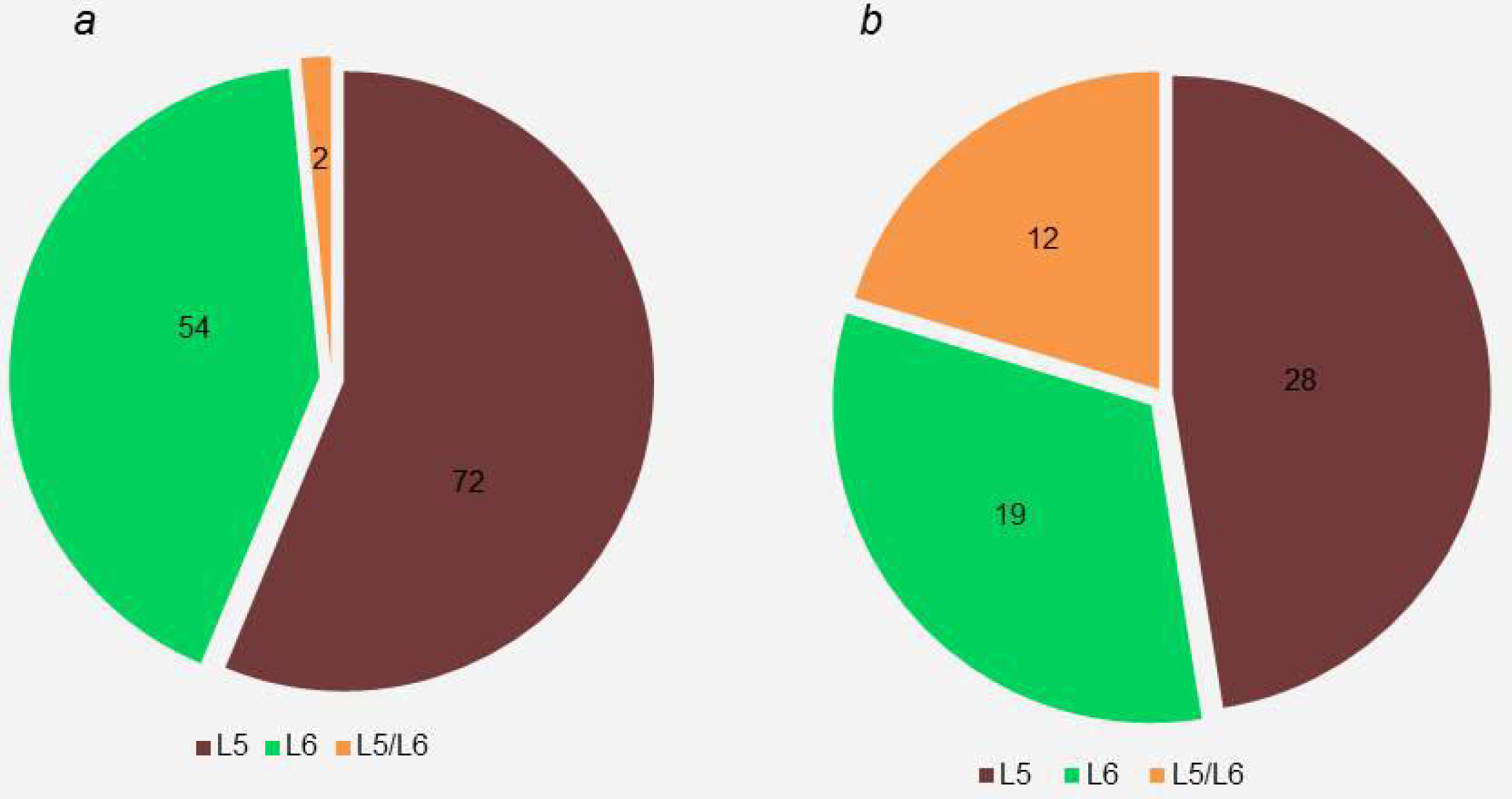
Number of human T cell epitopes (*a*) and human T cell antigens (*b*) with amino acid substitutions stratified by *Maf* lineage. Green represents L6-specifc mutant antigens or epitopes. Brown represents L5-specifc mutant antigens or epitopes. Yellow represents antigens or epitopes mutated in both L5 and L6 but at different loci with different amino acid substitutions.

### Conservation of human T cell epitopes of L5 is not affected by patient ethnicity

We previously reported an association between L5 and Ewe patient ethnicity^9,12^. Hence to test if conservation of T cell epitopes and/or the diversity of regulatory proteins in L5 was influenced by patient ethnicity, we estimated pairwise dN/dS for sequences encoding T cell epitopes and regulatory proteins of L5 genomes stratified by patient ethnicity (Fig 7). The median dN/dS of T cell epitopes were all below 1.0 irrespective of patient ethnicity (Fig 7A). However, the median dN/dS of regulatory proteins were marginally above 1.0 among L5 from Ewe TB patients and below 1.0 among L5 from non-Ewe TB patients (Fig 7B). There was no statistically significant difference between the estimated dN/dS of either the sequences encoding T cell epitopes (Fig 7A) or regulatory proteins (Fig 7B) between L5 from TB patients of Ewe and non-Ewe ethnicities. In addition, there was no difference in either the number of T cell epitopes with amino acid substitutions (Supplementary figure S6A) or the number of non-redundant SNPs (Supplementary figure S6B) between L5 strains from patients of the Ewe ethnicity and those of other ethnicities.

**Figure 7:**
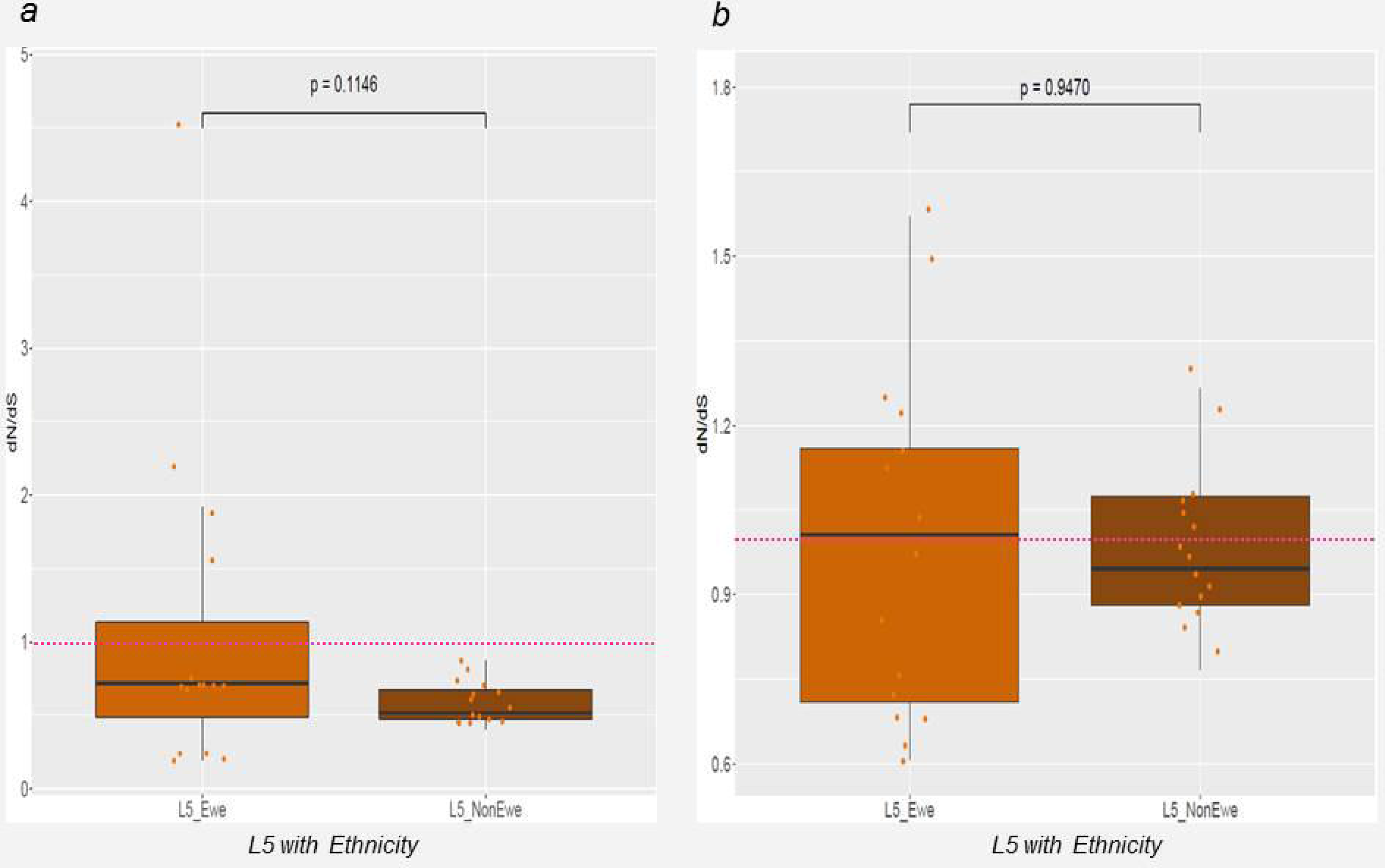
Pairwise dN/dS of sequences encoding human T cell epitopes (*a*) and genes encoding regulatory proteins (*b*) of L5 by patient ethnicity. L5 genomes from strains isolated from patients of the Ewe ethnicity (15 genomes) against, average values of 10 random samples of size 15 of L5 genomes of isolates from Non-Ewe patients.

## Discussion

In this study, we compared the largest collection of *Maf* genomes including both L5 and L6 reported so far. We found that, 1) at the whole genome level, L6 had significantly more pairwise nucleotide diversity, higher number of fixed SNPs as well as lower average pairwise SNPs relative to L5, 2) L5 had overall more conserved human T cell epitopes compared to L6, 3) conservation of T cell epitopes in L5 was not influenced by patient ethnicity, and 4) genes encoding regulatory proteins of L5 had lower pairwise nucleotide diversity but a higher ratio of non-synonymous to synonymous substitution rate than L6.

Our finding that *Maf* L6 has a higher number of fixed SNPs and higher average pairwise SNPs relative to L5 suggests that L6 has diversified more compared to L5 since the emergence of the two lineages^5,28,29^. This observation is corroborated by the higher genome-wide nucleotide diversity of L6 compared to L5. The higher diversity of L6 might be linked to an earlier emergence. However, recent whole genome-based phytogenies rooted on *Mycobacterium canettii* show that following the branch leading to *Maf* and all animal-adapted members of the MTBC defined by the characteristic deletion in RD9 ^30,31^, L5 branches off much earlier than L6^32^. Hence, L5 is ancestral to L6, a notion which is also supported by genomic deletion analyses which show that in addition to RD9, L6 and all the animal-adapted members of the MTBC harbor the deletions of RD7, RD8 and RD10^30^. Hence other factors are likely to account for the higher diversity of L6 compared to L5 in Ghana.

*Maf* is highly restricted to West Africa, and thus could be seen as an ecological specialist compared to the other MTBC lineages. Specialists are expected to harbour less diversity across strains compared to generalists^8^. The observed lower genome-wide nucleotide diversity of L5 hence supports the hypothesis that L5 might be a specialist maintained in West Africa by adaptation to specific human genotypes. In contrast, the higher genome-wide diversity of L6 indicates a generalist pathogen, and hence would have been expected to be globally distributed instead of displaying restriction to West Africa^4,8^. The observed diversity of L6 therefore may indicate a pathogen with a wider host range, supporting the hypothesis of maintenance in West Africa by possible environmental or zoonotic reservoir(s). Alternatively, the higher diversity of L6 coupled with the higher number of fixed SNPs could mean that it has a higher intrinsic mutation rate compared to L5.

Even though 90% of T cell epitopes were highly conserved in both L5 and L6 (Fig 5), which is in line with previous reports for the whole MTBC^26,8,33^, we found T cell epitopes in L5 to exhibit less nucleotide diversity and to be under purifying selection compared to L6. The purifying selection of mutations within L5 is comparable to that reported for the specialist sub-lineages of L4^8^. Interestingly, dN/dS within essential genes for survival in macrophages did not differ between L5 and L6 (Supplementary Figure S3) supporting the notion that the genes in this category perform key functions in both L5 and L6. Since T cell response might partially drive the pathogenesis of TB^34^, the relative conservation of T cell epitopes in L5 indicate that it might elicit a more efficient T cell response compared to L6 in its particular host population. This therefore suggests L5 may be a more human-specific pathogen and L6, with significantly more diverse T cell epitopes, a potential opportunistic environmental or zoonotic pathogen. Even though the conserved T cell epitopes of L5 could account for geographical restriction to West Africa and the association with the Ewe ethnicity^1,4,9,12^, we found no difference between the diversity of L5 isolated from TB patients of Ewe and those of non-Ewe ethnic backgrounds. The limited number of L5 genomes from Ewe TB patients could possibly account for the lack of observed difference in diversity of T cell epitopes of L5 from TB patients of Ewe and non-Ewe ethnicities, and hence larger sample sizes are required to explore this further. L5 isolated from TB patients of the Ewe ethnicity were shown to be distributed all across the L5 clade of the *Maf* phylogeny instead of clustering in a particular sub-clade (Supplementary figure S7). This suggests that, if L5 is indeed maintained in West Africa by its coevolution/adaptation with the Ewe ethnic group of West Africa (Cote d’Ivoire, Ghana, Nigeria, Togo and Benin)^9,12^, there is no specific sub-group of L5 that is responsible for this association but rather the whole of L5.

Members of the MTBC survive in the host mostly by modulation of the host immune response via the action of secretory proteins which form part of regulons controlled by specific regulatory proteins^35,36^. In addition, some regulatory proteins are involved in the regulation of transcription and translation of these secretory effectors as well as gene expression of other proteins involved with diverse functions^36,37^. Regulatory proteins in the MTBC hence play an important role in the survival and propagation of the bacteria. Therefore, our finding that regulatory proteins in L5 are under neutral selection (pairwise dN/dS = 1.03) compared to L6 in which they appear under purifying selection indicates that the mutations within regulatory proteins might be lineage-specific. This result is comparable to an earlier report comparing mutations within regulatory proteins between *Mtbss* and *M. bovis*, which found most of the *M. bovis* to harbor majority of the mutations^36^. As mutations within some regulatory proteins have been associated with attenuated virulence^38–40^, our observation could account for the reported attenuated virulence of *Maf* relative to *Mtbss*^14,15,17,41^. This calls for further comparative studies of regulatory proteins between L5, L6 and other MTBC lineages to ascertain the role of regulatory proteins.

Our data is limited by the fact that, the number of L5 genomes was almost 3 times the number of L6 genomes; however, we used 1,000x bootstrap sampling with replacement of both L5 and L6 of equal sample size to limit any possible bias when comparing both lineages due to differences in sample size. In addition, a number of the L5 genomes did not have data on ethnicity and hence affected the number of L5 isolated from patients of the Ewe ethnicity for which we used average estimates of 10 random samples of L5 isolated from patients of non-Ewe origin in comparisons to account for the different sample sizes.

In conclusion, our findings indicate that the two *Maf* lineages L5 and L6 are distinct in terms of genomic diversity, and selection pressure on T cell epitopes and regulatory proteins, possibly reflecting different ecological niches. Whereas L5 may be maintained in West Africa by its coevolution or adaptation with native West Africans, L6 may be maintained by an environmental reservoir, possibly a zoonotic source. This genomic analysis of *Maf* from Ghana gives a glimpse of the often neglected diversity within *Maf* and the MTBC overall. More studies are needed from representative genomes of *Maf* from across West Africa to understand the full diversity of these members of the MTBC. Improved knowledge of *Maf* will have implications for our understanding of human TB and the development of better control tools.

**Table 1:**
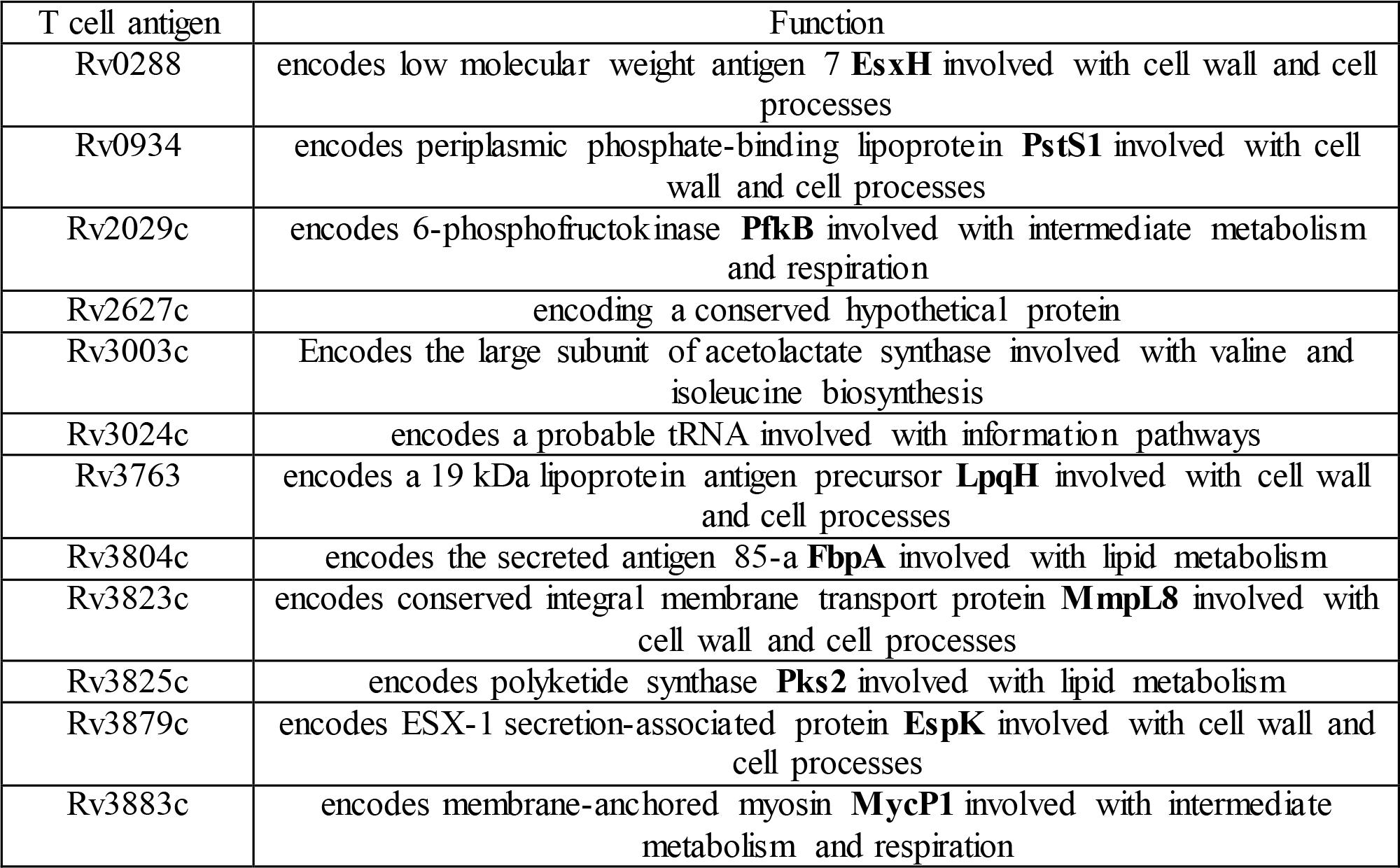
Functions of the 12 T cell antigens mutated in both L5 and L6.

## Acknowledgements

Bacterial Isolation and DNA preparations were done in the Biosafety level 3 facility at the Noguchi Memorial Institute for Medical Research, University of Ghana. Bioinformatics analyses were performed using the scientific computing core (sciCORE) at the University of Basel and the computing facility of the Wellcome Trust Sanger Institute, Genome Campus, Cambridge University. This work was supported by the Wellcome Trust Intermediate Fellowship awarded to DYM (Grant Number 097134/Z/11/Z) and by the Swiss National Science Foundation (grants 310030_166687, IZRJZ3_164171 and IZLSZ3_170834), the European Research Council (309540- EVODRTB) and SystemsX.ch.

### Author contributions

Conceived the idea: DYM, SG
Designed experiments: DYM, SG, IDO, MC, SRH, JP
Contributed reagents and performed experiments: IDO, MC, AAP, LSB, MC, SOW, AF, CL, GAA, AIY, AB, SYA, PA, CL, DB, SB, FG, PB, TK, SN, MA, SN, CB, BCDJ, JP and SRH
Analysed Data: IDO, MDC, SRH, LSB, SG, DYM
Wrote manuscript: IDO, MC, SG and DYM

All authors critically reviewed the manuscript

### Competing financial interests

None declared.

## Data availability

All the analyzed and/or generated data in this study are included in this article and its supplementary information files. Whole genome sequence reads have been submitted to the EMBL-EBI European Nucleotide Archive (ENA) Sequence Read Archive (SRA) with accession numbers provided in the supplementary document attached (Supplementary data S8).

## Supplementary Information

**Supplementary figure S1:**
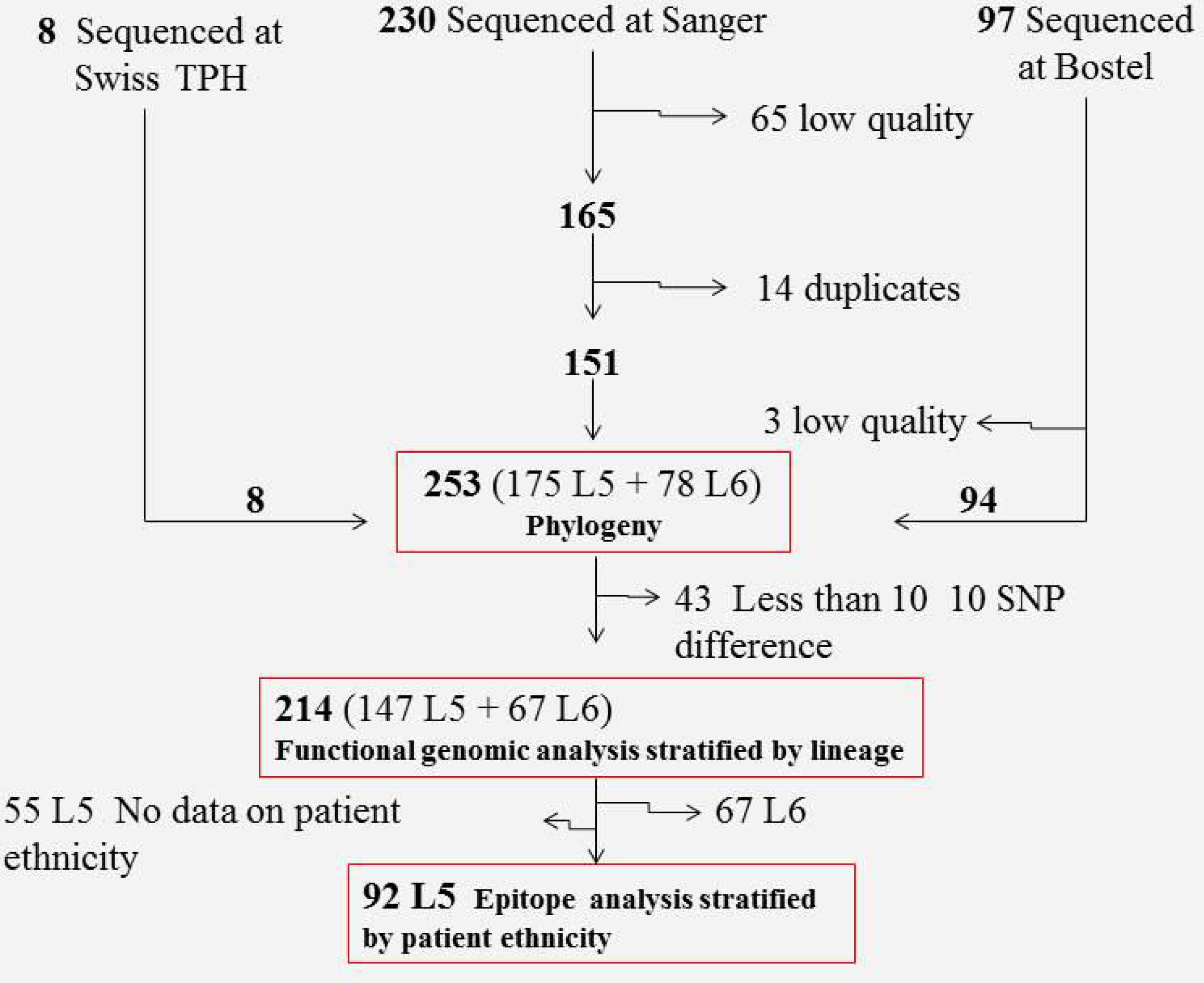
Flow chart of Ghanaian *Maf* genomes used for the study. Sites of DNA sequencing as well as number of genomes used for each specific analysis are indicated.

**Supplementary figure S2:**
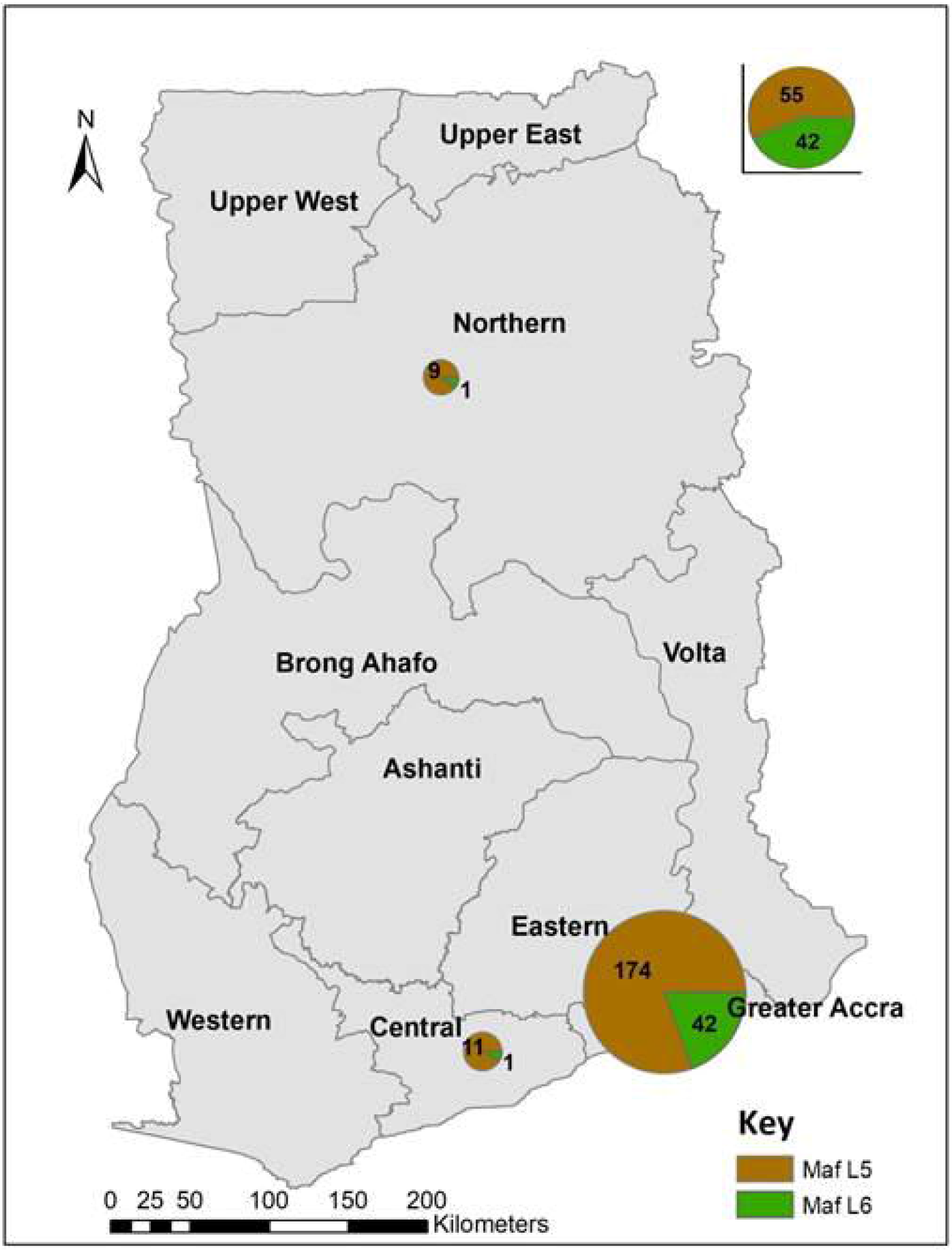
Map of Ghana showing the regional distribution of *Maf* isolates. The sizes of pie charts correspond to number of isolates from the respective regions.

**Supplementary figure S3:**
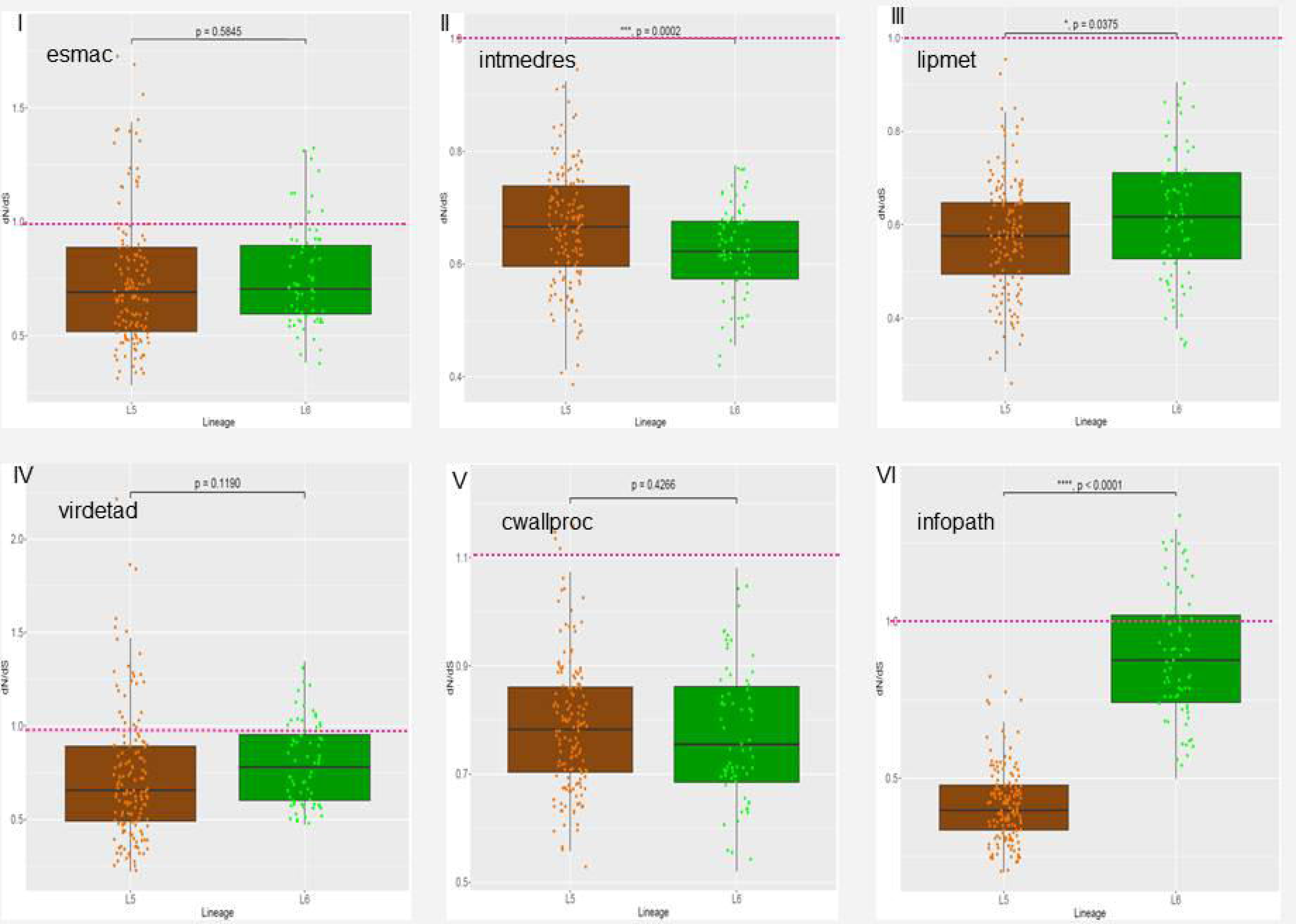
Pairwise dN/dS of genes of six functional categories using all the 147 L5 against the 67 L6 genomes. *esmac* - genes essential for growth in macrophages, *intmedres* - genes involved with intermediate metabolism and respiration, *lipmet* - genes involved with lipid metabolism, *virdetad* - genes involved with virulence, detoxification and adaptation, *cwallproc* - genes involved with cell wall and cell processes and *infopath* - genes involved with information pathways

**Supplementary figure S4:**
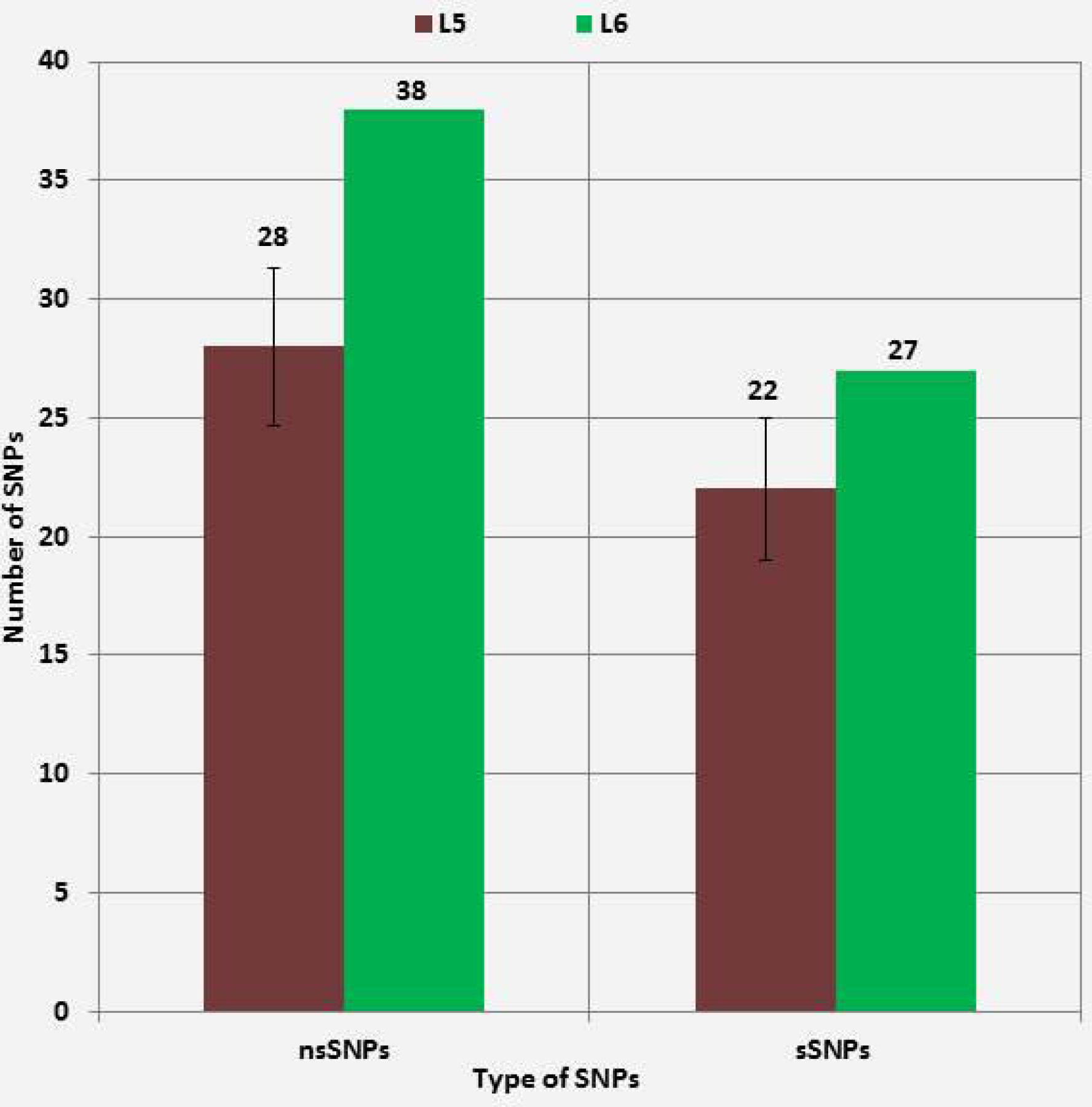
Number of non-redundant SNPs within human T cell epitopes stratified by *Maf* lineage. No difference (X-squared = 0.0055391, df = 1, p-value = 0.9407) between the proportion of nsSNPs between L6 (67 genomes) and L5 (mean values of 10 random samples of size = 67 with replacement)

Supplementary table S5: Mutated Human T cell antigens and epitopes of *Maf* with mutations stratified by lineage

**Supplementary figure S6:**
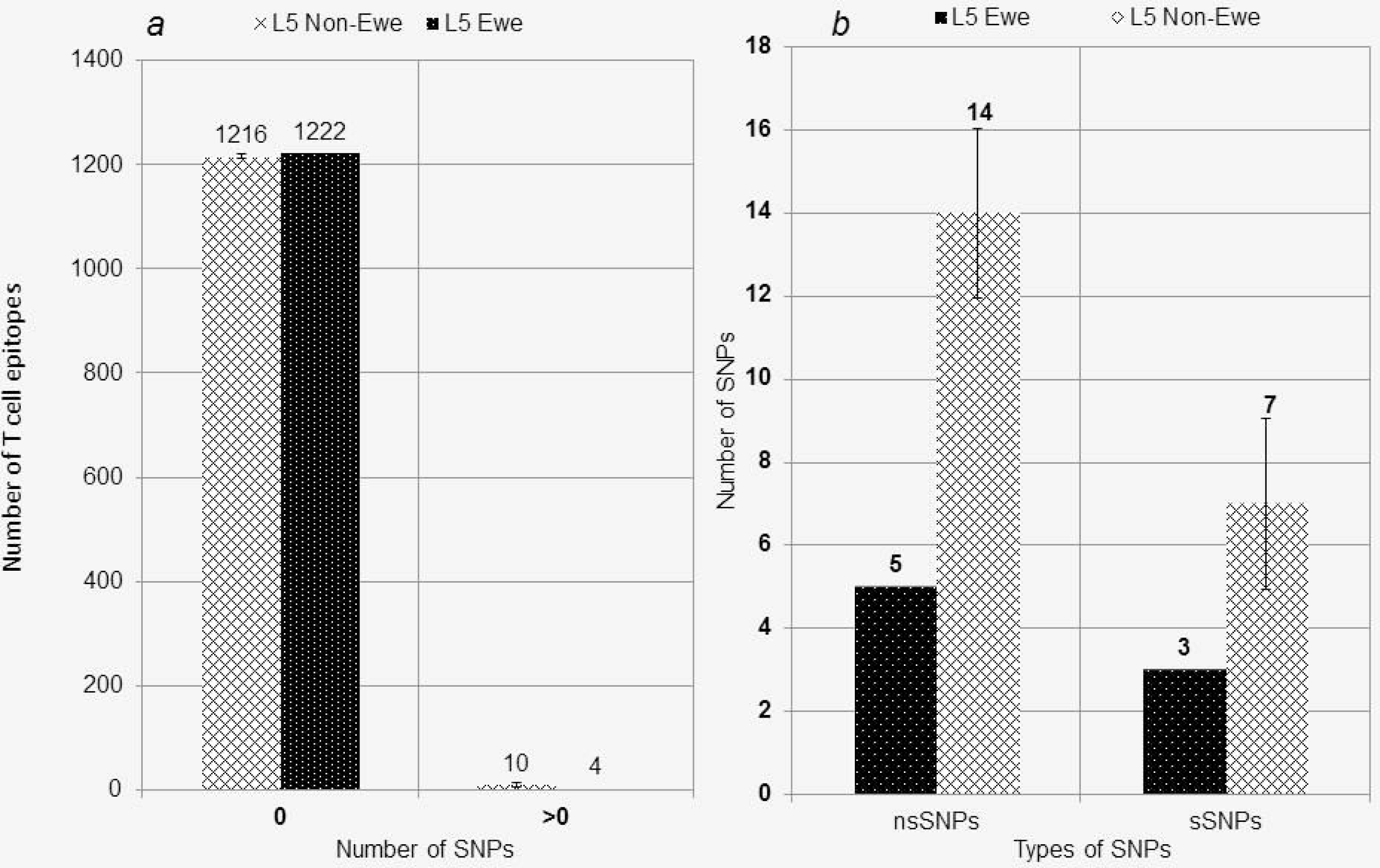
Number of epitopes with and without nsSNPs stratified by Lineage 5 with patient ethnicity (*a*). Number of non-redundant SNPs in epitopes stratified by ethnicity of L5 infected patients (*b*). (The values for the non-Ewe associated L5 are mean estimates of 10 random samples of size 15 with replacement)

**Supplementary figure S7:**
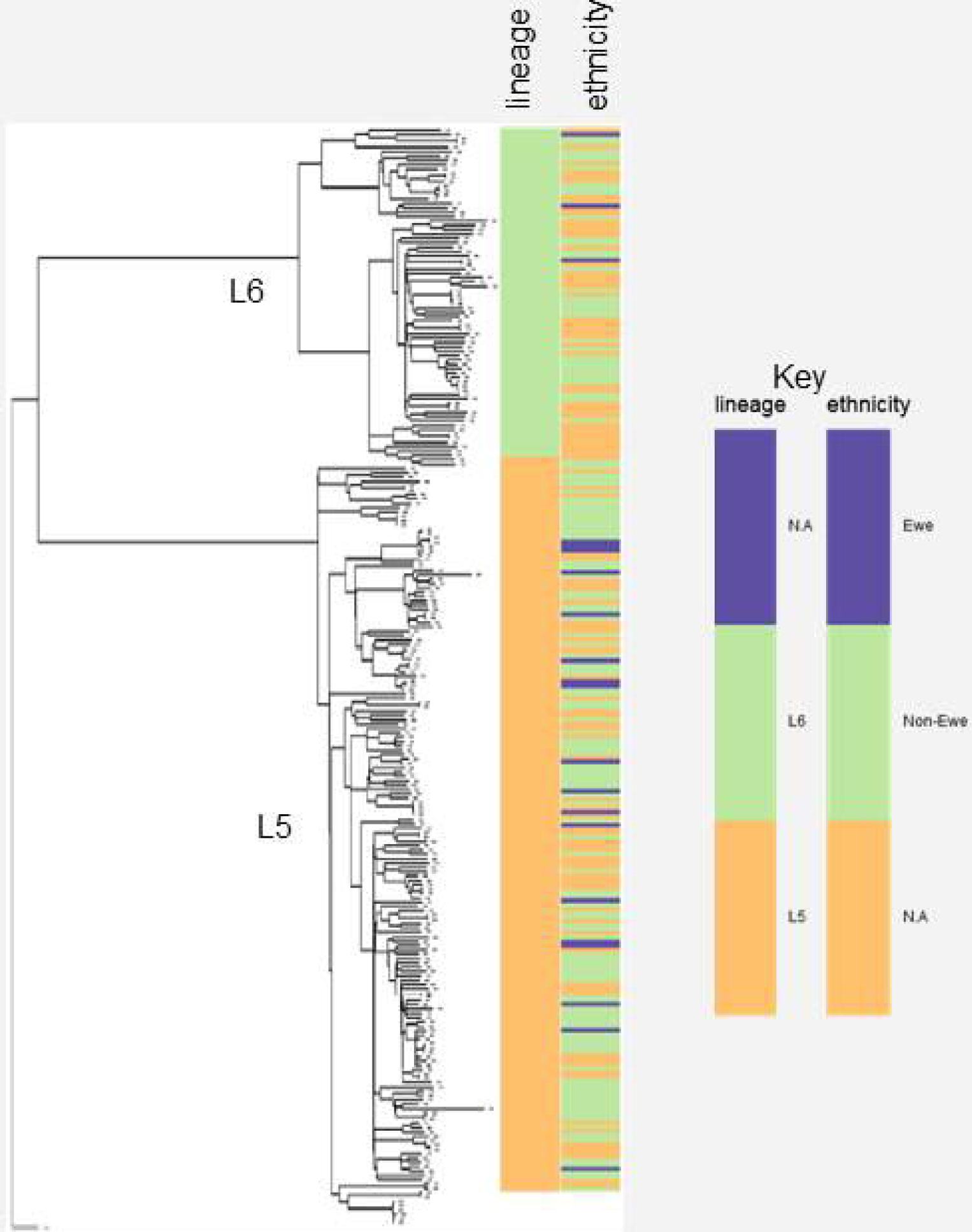
Uniform distribution ofL5 isolated from ethnic Ewe TB patients among the L5 clade.

Supplementary data S8: Genomes and accession numbers of*Maf* from Ghana.

## Methods

### Ethical Statement and Participant Enrollment

The study and its protocols were reviewed by the Scientific and Technical Committee and approved by the Institutional Review Board (IRB) of the Noguchi Memorial Institute for Medical Research, Legon-Ghana with Federal Wide Assurance number FWA00001824.

### *Mycobacterium africanum* Strains

Isolates used for this study were cultivated from July 2007 to November 2014 in Ghana^9,12^, West Africa, involving consecutive sputum smear positive pulmonary TB cases recruited from two different studies with isolates spanning four regions of Ghana (three in the south and one from the North) (supplementary figure S2).

### Mycobacterial Sub-Culturing and Chromosomal DNA Extraction

*Mycobacterium africanum* strains were revived by sub-culturing on Lowenstein Jensen (LJ) slants; one supplemented with 0.4% sodium pyruvate the other with glycerol to enhance the growth of Lineage 5 and Lineage 6 strains of respectively. The cultures were incubated at 37 °C and monitored regularly until growth was observed. When confluence was achieved, five loops full of colonies were fetched into 2 mL cryo-vials containing 1 mL of sterile nuclease-free water, heat-inactivated at 98 °C for 60 minutes for DNA extraction using a hybrid DNA extraction protocol^42^. The isolates were confirmed MTBC by PCR amplification of IS *6110* genotyped as *Maf* by large sequence polymorphism (LSPs) detecting region of difference (RD) 9 and 12^43^. Lineage identification was achieved by spoligotyping as previously described^44^. Strains confirmed as belonging either L5 or L6 were sequenced by the illumina platform at the Wellcome Trust Sanger Institute, United Kingdom.

### DNA Sequencing, Mapping of Sequence Reads, Variance Calling and Generation of Whole Genome Fasta files

Samples were sequenced as multiplexed libraries on the Illumina HiSeq platform to produce paired end reads of 125 nt in length. Genomes provided by the Research Center Borstel was obtained by sequencing DNA libraries prepared with the Nextera XT kit and run on Illumina MiSeq (250 and 300 bp, paired end) and NextSeq (150 bp, paired end) according to the manufacturer’s instruction (Illumina, San Diego, USA). The FastQ files containing the raw paired-end reads were processed using a python pipeline developed in house as follows. The reads were first adapter-and quality-trimmed with Trimmomatic v0.33^45^. Reads lower than 20 bp were not kept for the downstream analysis. Overlapping paired-end reads were then merged with SeqPrep (https://gthub.comjstjohn/SeqPrep). The resulting filtered reads were mapped to a hypothetical reconstructed MTBC ancestor^26^ with BWA v0.7.12^46^. Duplicated reads were marked by the MarkDuplicates module of Picard v 2.1.1 (https://sithub.com/broadinstitute/picard). The RealignerTargetCreator and IndelRealigner modules of GATK v.3.4.0 (https://software.broadinstitute.org/gatk/download/archive) were used to perform local realignment of reads around indels. SNPs were called with Samtools v1.2 (https://sourceforge.net/projects/samtools/fles/samtools/1.2/) and VarScan v2.4.1^47^ using the following thresholds: minimum mapping quality of 20, minimum base quality at a position of 20 and minimum read depth at a position of 7X. SNPs were considered fixed at a frequency of ≥ 90% and alleles were considered ancestral when the SNP frequency was ≤ 10%. Furthermore, SNPs were called only if the alternative basecall was supported by at least five reads and without strand bias. All variants were annotated using snpEff v4.11^48^, in accordance with the *M. tuberculosis* H37Rv reference annotation (AL123456.3). SNPs falling in regions with at least 50 bp identity to other regions in the genome were excluded from the analysis.

### Generation of Variable Positions and Phylogenetic Analysis

The variable SNPs alignment was obtained by concatenating the SNP calls present in the variant calling file of each genome, using the IUPAC nucleotide ambiguity codes for heterozygous calls. A position was considered variable if at least one genome had a SNP at that position. Called deletions and positions not called according to the minimum threshold of 7 were encoded as gaps. Positions for which the proportion of gaps exceeded 50% were excluded from the alignment. Maximum likelihood phylogeny of the variable positions with 1000 bootstraps was then generated using RAxML version 8.2.3^49^ with GTR substitution matrix and other default settings with the final tree evaluated and optimized under GAMMA with accuracy of 0.1 Log likelihood units. The best tree was then, rooted on *M. canettii* and annotated using figtree (http://www.webcitation.org/getfile?fileid=27177ee8dd2f34cfd254b9c5e6c6fdf4b65329f6).

### Comparative genomics analysis of isolates using genes encoding proteins of 8 functional categories

Experimentally confirmed human MTBC T cell epitope (1,226 epitopes) sequences (spanning 304 antigens with some overlapping sequences) retrieved from the Immune Epitope Database (IEDB), tested in human T cell assays, with no major histocompatibility complex (MHC) restrictions and have genomic coordinates in the H37Rv reference strain^32,8^ were *in silico* extracted from the fasta whole genome files and concatenated excluding sequence redundancy using customized bash algorithms. Complementary sequences of epitopes encoded by the reversed strand were first transcribed before the concatenation to have all the sequences in the same direction. In addition, MTBC genes of other seven functional categories namely those encoding regulatory proteins (regprot; 196), genes involved with lipid metabolism (limpet; 267), genes involved with intermediate metabolism (intmedres; 917), genes involved with virulence, detoxification and adaptation (virdetad; 216), genes involved with information pathways (infopath; 234), genes involved with cell wall and cell processes (cwallproc; 768) and genes essential for growth in macrophages (esmac; 125) according to the tuberculist database^50^ were also retrieved and concatenated as described above excluding genes involved with drug resistance.

### Estimation of Pairwise Nucleotide Diversity

Pairwise SNP distances of the whole genome excluding sites associated with drug resistance, concatenates of T cell epitopes and the genes of other seven functional categories were calculated with the *dna.dist* function of *ape* package^51^ of R version 3.2.3^52^ as previously described^8^. Average pairwise nucleotide diversity per site (*π*) and confidence intervals for the *π* was calculated as previously described^8^ and plotted with *ggplot2* package implemented in R. The upper and lower levels of confidence were attained by estimating the 97.5^th^ and 2.5^th^ quantiles of the *π* distribution obtained by bootstrapping (1000 replicates) as previously described^8^. Non-overlapping confidence intervals of *π* were taken as evidence of statistically significant differences^53,54^. Details of the algorithm for this analysis are available upon request.

### Estimation of Pairwise dN/dS

The concatenates of the human T cell epitopes and the other genes of seven functional categories were also used for estimation of dN/dS ratios stratified by lineage. As a follow up, dN/dS of T cell epitopes and regulatory proteins were also estimated for 15 L5 genomes from Ewe TB patients and 77 from non-Ewe TB patients. The dN/dS estimates were calculated with all polymorphic sites within each lineage using the *kaks* function of the *seqinr* packgage^55^ as previously described^8^ and box plotted using *ggplot2* package in in R version 3.2.3. Statistical difference of the estimates between the *Maf* lineages was accessed using the non-parametric Wilcoxon rank-sum tests with continuity correction in R version 3.4.0.

### Human T cell Epitopes with Non-Synonymous SNPs and Count of Non-Redundant SNPs

Synonymous and non-synonymous mutations within the coordinates of each epitope were extracted from the variant calling file (VCF) obtained for each genome. The specific human T cell epitopes with non-synonymous SNPs were compared between the *Maf* lineages for lineage-specific mutated epitopes and *Maf*-specific mutated epitopes. furthermore, the number of pairwise non-redundant SNPs was estimated for the *Maf* lineages (67 L6 genomes and the 10 random samples of L5 of equal size as L6) as well as L5 genomes stratified)y patient ethnicity (15 L5 from Ewe patients and 10 random samples of L5 from non-Ewe TB)atients of size 15) using Mega6^56^. The number of SNPs per each group was plotted and compared)between the groups using the fisher’s exact test for statistical significance in R version 3.2.3.

